# Prevalence of Class 1 integron in disease-causing carbapenem-resistant *Escherichia coli* in a tertiary hospital in Kathmandu, Nepal

**DOI:** 10.1101/2024.02.28.582515

**Authors:** Suchitra Thapa, Basudha Shrestha, Manisha Shrestha, Yojana Pokhrel, Reshma Tuladhar, Elita Jauneikaite, Dev Raj Joshi

## Abstract

**Background:** High prevalence of carbapenem resistance genes have been abundantly reported within clinical *E. coli* in Nepal but their co-prevalence with class-1 integron is poorly documented. These Class I Integrons play pivotal role in evolution and spread of antimicrobial resistance (AMR). Therefore, it is necessary to highlight their epidemiological significance, so this study determined the prevalence of Class 1 integrons within Carbapenem resistance *Escherichia coli* (CREc) population.

**Methods:** Fifty-two clinical *E. coli* with confirmed resistance to Meropenem were collected between January 2020–April 2022 in Kathmandu Model Hospital, and undergone antimicrobial susceptibility testing for 11 antimicrobial classes. Carbapenemase production was confirmed by eCIM and mCIM methods. Presence of Class I integron and carbapenemase genes, *bla*_NDM-1_, *bla*_KPC_, *bla*_IMP_, *bla*_VIM_, *bla*_OXA-23_, *bla*_OXA-48_, was confirmed by PCR method.

**Results:** The result revealed 92.3% (n=48/52) of the isolates harbored *intI1* gene. Among integron-positive population, 97.9% (n=47/48) were carbapenemase producers which were further identified as metallo β-lactamase (MBL) producers (82.9%, n=39/47) and serine β-lactamase (SBL) producer (17%, n=8/47). Carbapenemase gene *bla*_NDM-1_ (79.2%, n=38/48) was highly prevalent within integron-positive isolates, followed by *bla*_OXA-23_ (62.5%, n=30/48) and *bla*_OXA-48_ (54.2%, n=26/48). All integron-positive CREc isolates were resistant to minimum 16 antibiotics tested, but were susceptible to Colistin, Polymixin B, and Tigecycline.

**Conclusion:** In conclusion, Class 1 integrons were highly prevalent within clinical CREc in Nepal, and their co-prevalence with other CRE gene is concerning due to possibility of wider spread. Therefore, an in-depth genomic analysis of Class 1 integrons is needed to understand the integron mediated AMR in high risk clones like CREc and inform intervention measures to stop spread of such pathogens.

## Introduction

Carbapenems, a broad-spectrum antibiotic, are the last resort treatment drug for many complicated infections. Resistance to carbapenem has been emerging and increasing globally, becoming a public health threat.^1^ The most common mechanism of carbapenem resistance is acquisition of resistance genes, particularly the carbapenemase enzyme producing trait. Genes encoding these carbapenemase are usually found in mobile genetic elements and are easily transferable by plasmids. ^2-3^

Integrons are often found inserted into mobile plasmids and are responsible for the dissemination of many different cassette-associated antibiotic resistance genes ^4^ including CRE genes. They are considered as drivers of antimicrobial resistance ^5^ due to their ability to capture and integrate antibiotic resistance genes, thus forming novel antimicrobial resistance cassettes particularly in Gram-negative clinical isolates.^6^ So considerably they play an important role in both dissemination and evolution of antibiotic resistant genes. So far, five classes of integrons (class 1, 2, 3, 4 and 5) have been identified based on the integrase gene (*intI*) sequence, but only three classes (class 1, 2 and 3) are known to be responsible for multi-drug resistance (MDR). ^5,7^ And among the three classes, Class 1 integrons are the most prevalent in clinical isolates and carry single or multiple gene cassettes.^4,7^ Reportedly, these class 1 integrons are widely distributed in clinical and environmental Gram-negative bacteria.^4^ Some studies have even reported 100% prevalence within MDR bacteria ^7-10^ which were mostly *E. coli*.

In Nepal, few studies have reported the occurrence of Class 1 Integron in both clinical^11-12^ as well as environmental isolates^13-15^; which were mostly multiple antibiotic resistant isolates including MDR *Salmonella* spp ^12^ and extended-spectrum β-lactamase producing (ESBL) *E. coli*.^11^ Class 1 integron was reported only in bacteria of clinical origin ^12^ when compared to environmental isolates in Nepal. Despite the reported dominance in clinical setting and continuous changing trend of antibiotic resistance, there is no recent information on class 1 integrons from high risk clones like CRE from Nepal. Therefore, considering the vital role of class 1 integrons in accelerating AMR evolution and dissemination,^16^ it is important to know the epidemiology of integron-mediated antibiotic resistance in clinical isolates particularly the multidrug-resistant clones. Therefore, this study aimed at determining the prevalence of class 1 integrons as well as their co-prevalence with other CRE genes in CREc of clinical origin. And by determining the prevalence of Class 1 integron along with other CRE genes in clinical settings, we can identify the risk clones early on and work towards placing appropriate interventions to reduce the dissemination of such risk clones within community and clinical settings.

## Materials and Methods

### Sample and bacterial strains

Fifty-two *E. coli* with confirmed resistance to Meropenem (tested as part of routine diagnostics procedures using disc diffusion method following CLSI guidelines^17^) were collected during January 2020–April 2022 in Kathmandu Model Hospital, Kathmandu, Nepal. *E. coli* were isolated following microbiology laboratory guidelines^18^ from different clinical samples from inpatients, outpatients, and referral cases received at the Microbiology Unit of the hospital. *E. coli* isolates were preserved by mixing a heavy inoculum of overnight culture from Nutrient agar plates in tryptic soy broth with 20% glycerol and freezing at -80 °C.

### Antimicrobial Susceptibility Testing

The antimicrobial susceptibility testing was done using the Kirby Bauer disc diffusion method following CLSI guidelines.^17^ Twenty-four different antibiotic discs (Mast Diagnostics, Germany) representing eleven different antibiotic classes were used: Aminoglycosides (Amikacin 30μg, Gentamycin 10μg), Cephalosporins (Cefotaxime 30μg; Ceftazidime 30μg; Cefepime 30μg; Cefoxitin 30μg), Penicillins (Amoxicillin 25μg; Piperacillin 100μg), Penicillin/β lactamase inhibitor combinations (Piperacillin-Tazobactam 110μg; Amoxicillin-Clavunate 30μg; Ampicillin-Sulbactam 10/10µg; Cefoperazone/Sulbactam 85μg), Phenicols (Chloramphenicol 30μg), Tetracyclines (Tigecycline 15μg; Doxycycline 30μg), Fluoroquinolones (Ciprofloxacin 5μg; Levofloxacin 5μg; Ofloxacin 5μg), carbapenems (Meropenem 10μg; Ertapenem 10μg), Nitrofurans (Nitrofurantoin 300μg) Lipopeptides (Polymixin B 300μg; Colistin 10μg) and Sulfonamides (Sulfamethoxazole-trimethoprim 25μg). Additionally, agar dilution method was carried out to determine minimum inhibitory concentration (MIC) for Meropenem (PHR1772-500MG, Pharmaceutical Secondary Standard, SIGMA-ALDRICH) following CLSI guideline (31^st^ edition 2021).^17^ Quality control was ascertained by using *E. coli* ATCC 25922 strain as control in both, disc diffusion and MIC experiments.

### Phenotypic detection of carbapenemase production

All isolates were subjected to phenotypic confirmation for carbapenemase production, which was done by the modified carbapenem inactivation (mCIM) and the EDTA-carbapenem inactivation (eCIM) methods, as described by CLSI guidelines (31^st^ edition 2021).^17^

### DNA extraction

Overnight cultures of single colony of each bacterial culture in LB broth were centrifuged twice, and the pellet was suspended in 100 µL of 50 mM NaOH. The suspension was consequently boiled in a boiling water bath for 5 minutes, cooled immediately at 4 °C, and then mixed with 16 µL of 1M Tris-HCl. Then it was centrifuged at 12,000 rpm for 5 minutes and equal volume (116 µL) of cold absolute ethanol was added before centrifugation again for 15 minutes as previously described.^19^ The pellet was then air dried and suspended in 50 µL of TE buffer. The suspension was used as a DNA template in PCR reactions. The purity and concentration of extracted DNA were determined through 260/280 nm absorbance measures using the Nano Drop spectrophotometer 1000 (Thermo Scientific).

### PCR amplification of carbapenemase genes and *intI1* gene

Amplification of the extracted DNA was performed for six β-lactamase genes of which three were Metallo β-lactamase (*bla*_NDM-1_, *bla*_KPC_, *bla*_IMP_) and three were Serine β-lactamase (*bla*_VIM_, *bla*_OXA-23_, *bla*_OXA-48_) and the *intI1* gene by PCR methods ^20-23^ (Supplementary tables S1 and S2) in a Thermocycler (BIO-RAD T100™) using specific primer sets (Supplementary table S2). The amplified products were run on 1.5% agarose gels containing 0.5 µg/mL of ethidium bromide in running buffer and visualized by a gel documentation system (azure biosystems, Version 1.5.0.0518). *E. coli* ATCC 25922 DNA was used for positive control and PCR mix without DNA template was used as negative control.

### Statistical Data Analysis

All data were entered in MS-Excel. Continuous data and categorical data were used for descriptive statistics. MIC50 (MIC value at which 50% isolates growth is inhibited), MIC90 (MIC value at which 90% isolates growth is inhibited) and MIC90/MIC50 (Dispersion value of MIC) was determined manually. The data analysis was done using IBM SPSS Statistics (Version 20), R software (version 4.3.2), and WHONET 2020 software (version 2020).

### Ethical approval and patient consent

The study has been approved by Institutional Review committee of the Kathmandu Model hospital and Nepal Health research council (Approval ID: 378/2021). Written-informed consent was taken from the patients whose sample was included in this study.

## Results

### Demographic distribution of CREc

A total of 52 carbapenem-resistant *E. coli* (CREc) were isolated as part of the study. Among total 51.9% (n=27/52) isolated were recovered from clinical specimens of male patients while 48.1% (n=25/52) isolates were from those of female patients (Supplementary table S3). The isolates were majorly recovered from urine sample (n= 28/52; 53.8%) followed by pus sample (n= 12/52; 23.1%). The majority of sample (n=16/52; 30.8%) belonged to either out-patient department or were referral cases (Supplementary table S3).

### Prevalence of Class 1 integrons within CREc

Based on the presence of *intI1* gene (Supplementary figure F1), a total of 48 (92%) CREc isolates were class 1 integron positive. Within the integron-positive population, higher percentage was observed in urine isolates (n=27/48; 56.2%) followed by the isolates from 60 above age group (n=19/48; 39.6%) and out-patient department (n =16/48; 33.3%) while no distinct difference was observed in sex category of patients (Supplementary tables S4 and S5).

### Antibiotic resistant profiles of CREc and Class 1 integron-positive isolates

All 52 isolates were resistant to Amoxicillin, Piperacillin, Ampicillin-sulbactam, Cefoperazone-sulbactam, Cefotaxime, Cefepime, Cefoxitin, Ceftazidime, Ciprofloxacin, Levofloxacin, Ofloxacin, Etrapenem and Meropenem and (Figure 1; Supplementary table S6). Five isolates (n=5/52; 9.6%) showed resistance against 21 antibiotics, out of 24 tested (Supplementary table S7). Within the integron-positive population, all the isolates (n=48/48; 100%) showed resistance against Penicillins (AMX, PI), Penicillins with inhibitor (AMS, CFS), Cephalosporins (CTX, CX, CAZ, CPM), Fluoroquinolones (CIP, OF, LE) and Carbapenem (MEM, ETP) (Supplementary table S6). On comparing the antibiotic resistant profile of each antibiotic within integron positive and negative population, no significant difference in distribution was observed except Meropenem (p =0.02).

**Figure 1:**
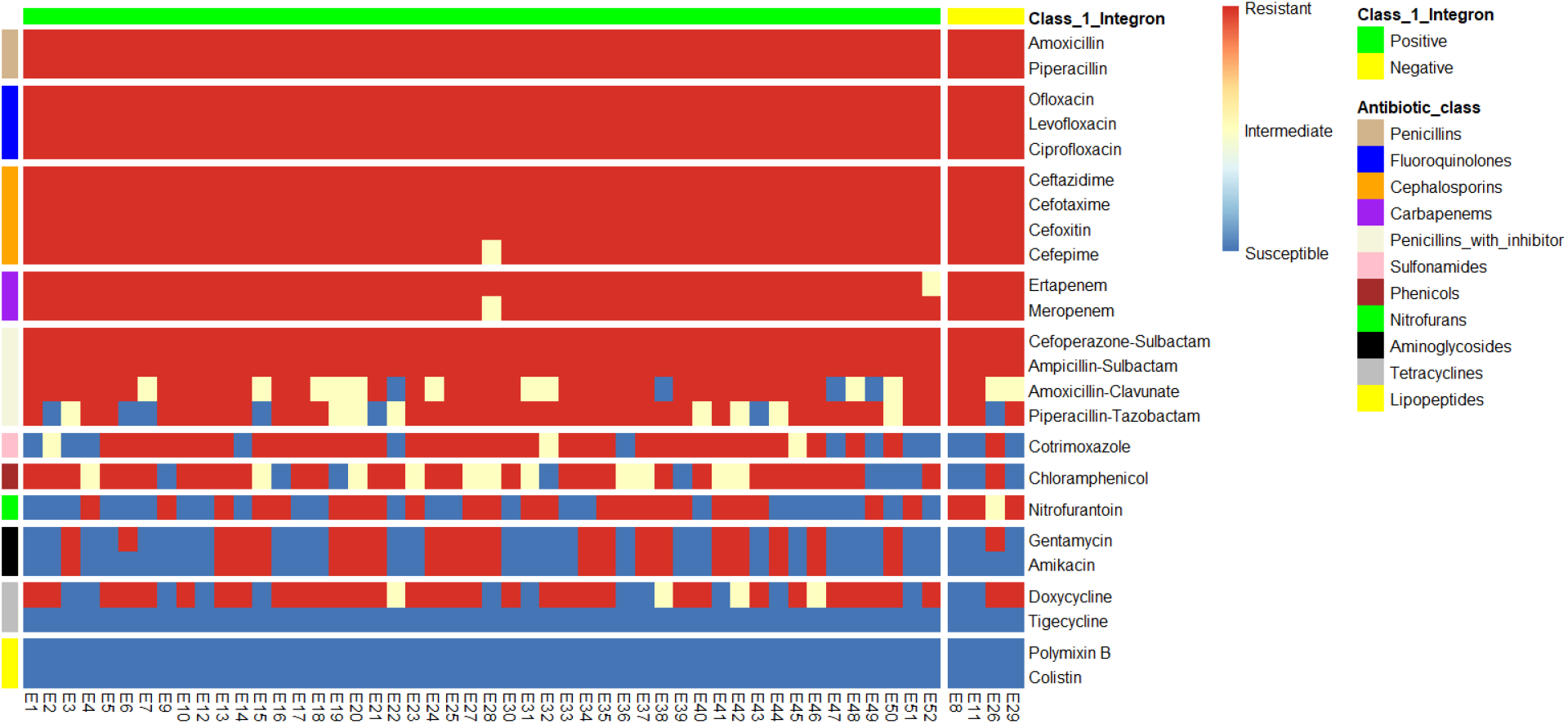
Heat map of antibiotic resistant profile of integron-positive and negative population against different classes of antibiotics.

Based on phenotypic antibiotic susceptibility testing results, all 52 isolates were considered as multi-drug resistant *E. coli* as most strains were resistant at least to 6 different class of antibiotics (Supplementary table S7). Within integron-positive population, three isolates (3/48; 6.3%) showed phenotypic resistance to 6 classes of antibiotics (Supplementary table S7) and seven isolates (7/48; 14.6%), showed resistance to 10 different classes of antibiotics out of 11 classes used in this study. The most common resistance profile combination (n=9/48; 18.8%) was Penicillins - Penicillins with inhibitor – Fluoroquinolones – Cephalosporins - Carbapenems-Tetracylines – Phenicols - Sulfonamide (Supplementary table S7). The four integron-negative CREc isolates showed diverse antibiotic resistance patterns as 2 isolates were resistant to 6 antibiotic class and 2 were resistant to 9 antibiotic class (Supplementary table S7).

### Meropenem MIC of CREc and Class 1 integron-positive isolates

The MIC of Meropenem for all 52 isolates ranged from 0.5 to 512 mg/L. And based on Meropenem CLSI breakpoint (S≤ 1mg/L and R ≥4 mg/L), 50 (96.2%) isolates were considered resistant to Meropenem while one as intermediate and one as susceptible (Supplementary table S8). The MIC_50_ and MIC_90_ were calculated to be 128 mg/L and 256 mg/L respectively and the ratio of MIC_90/50_ was 2. The MIC_50_ and MIC_90_ value indicates clustering far away from the cut-off value (R ≥4 mg/) within the Meropenem resistant isolates while the MIC_90/50_ value was low which suggests the dispersion of MIC from 50% isolates to 90% isolates was not large.

### Carbapenemase producers among CREc and Class 1 integron-positive isolates

Phenotypic confirmation test for carbapenemase production revealed that 51 (98.1%) were carbapenemase producers while only one (1.9%) was non-producer. Also, majority of carbapenemase producer were integron-positive (n= 47/51; 92.2%). Likewise, within the integron-positive population, higher proportion of carbapenemase producers were isolated from the patients of >60 years’ age group (n=18/48; 37.5%) and urine sample (n=26/48; 54.2%) (Supplementary table S4).

Further, based on the result of mCIM and eCIM (Supplementary Figure F2), 41 (78.85%) isolates were confirmed as MBL producer and 10 (19.23%) isolates were SBL producer.

Within integron positive population, MBL producers (n=39/48; 81.3%) were in greater proportion than SBL producers (n=8/48; 16.7%). Further, these MBL producers within integron positive population, were mostly isolated from patients of >60 years age group (n=18/48; 37.5%) and urine sample (n=23/48; 47.9%) (Supplementary table S4).

### Carbapenemase gene prevalence in CREc and Class 1 integron-positive isolates

Among the β-lactamase gene (Supplementary Figure F3), the prevalence percentage of NDM-1, OXA-23, OXA-48 and VIM encoding genes was found to be 75% (n=39/52), 65.4% (n=34/52), 53.4% (n=28/52) and 5.8% (n=3/52), respectively. While IMP and KPC were not detected in any of the isolates.

Majority of *bla*_NDM-1_ positive isolates were from patients with >60 years age group (n= 16/39; %), urine sample (n= 23/39; %) and out-patient department (n= 14/39; %). Likewise, for *bla*_OXA-23_, greater frequency was observed in patients with >60 years age group (n= 14/34; %), urine sample (n= 19/34; %) and out-patient department (n= 11/34; %). While for *bla*_OXA-48_, greater frequency was observed among patients with >60 years’ age group (n= 10/28; %), urine sample (n= 14/28; %) and referral hospital (n=10/28; %). For *bla*_VIM_, greater frequency was observed among 31-45 age group (n=2/3; %), urine sample (n= 3/3; 100%). and out-patient department (n=2/3; %)

Among the integron-positive population, the occurrence of NDM-1, OXA-23, OXA-48 and VIM encoding gene was 79.2% (38/48), 62.5% (30/48), 54.2% (26/48) and 6.2% (3/48), respectively (Supplementary table S4). Additionally, several combinations of carbapenemase encoding gene within integron-positive population was observed and the most predominant combination was NDM-1: OXA23 (n=12/48; 25%) followed by NDM-1: OXA-23: OXA-48 (n=11/48; 22.9%) and NDM-1: OXA-48 (n=7/48; 14.6%) (Figure 2)

**Figure 2:**
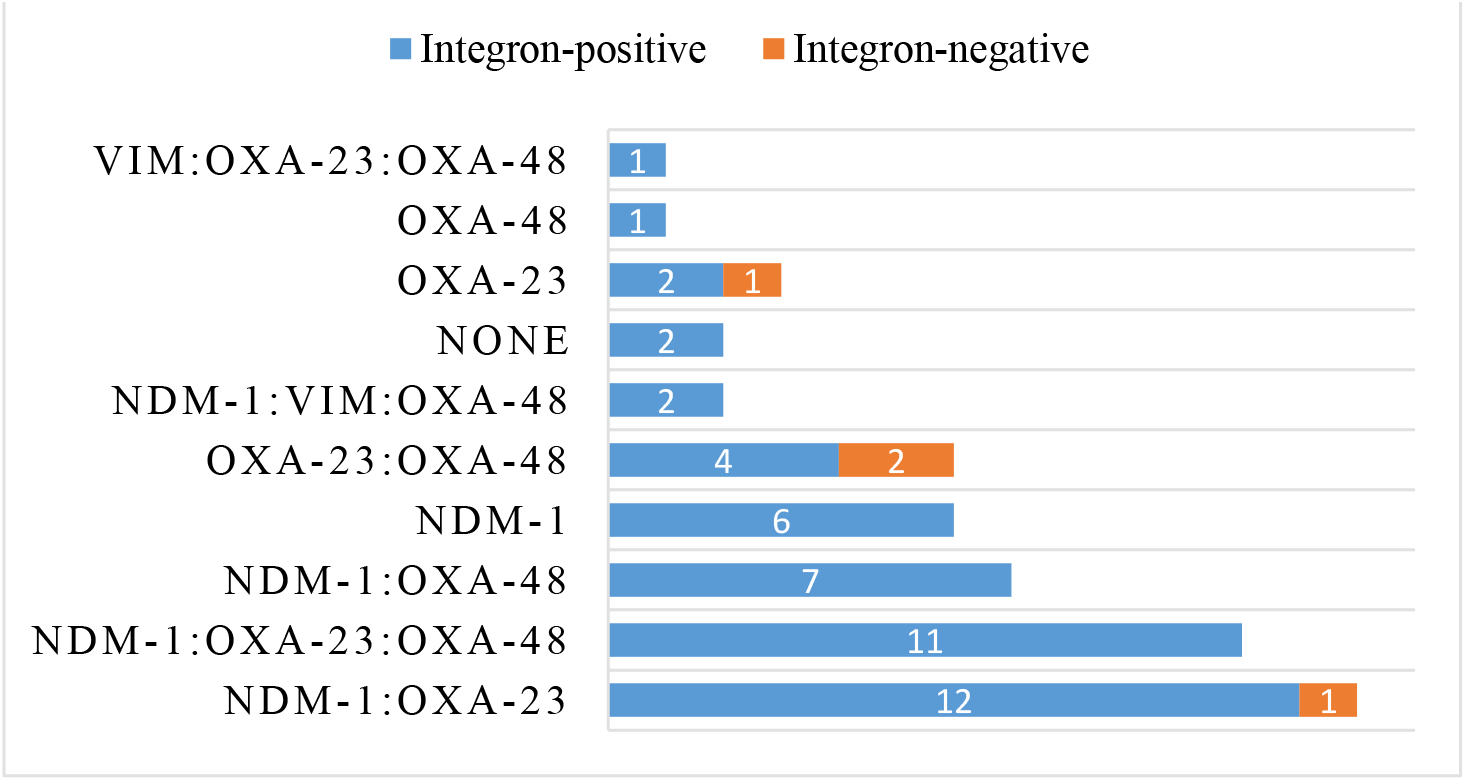
Frequency of different combination of four carbapenemase (NDM-1, OXA-23, OXA-48, VIM) encoding gene within Integron-positive (n=48) and negative population (n=4). Note: KPC and IMP were not detected.

## Discussion

Class 1 integrons are mobile genetic element that are involved in helping to transfer AMR genes via horizontal gene transfer and are important to help to understand the molecular epidemiology of clinically relevant and, commonly, multi-drug resistant bacterial pathogens. In this study, the prevalence of the Class-1 integron gene, *intI1*, among CREc isolates was found to be high – 92% (n=48/52). Though not similar, a varying prevalence percentage of class 1 integrons has been well documented within multidrug-resistant *E. coli* from humans,^10,23-29^ animals,^29-30^ and the environment.^7,13-15^ One possible reason for such high prevalence may be that integrons occur widely in Enterobacteriaceae family, and are strongly correlated with MDR.^8-9,27-28,30^ Also, their ability to freely move within and between species with the help of conjugative plasmids may also be a contributing factor.^16^ Besides, their increased occurrence in pathogenic *E. coli* over the period of 100 years is already well documented.^31^

As for demographic distribution (Supplementary table S4), majority of the integron-positive isolates were among the 60 and older age group, followed by the 31–45 years old age group. The vulnerable health condition and many such predisposing conditions can be contributing factors to the higher prevalence among the 60 and older age group.^32^ Besides, a study on uropathogens and age categories has revealed that antibiotic resistance occurrence increase with age.^28,32^

In the case of the department, out-patient department samples showed greater prevalence, followed by ward samples. This suggests that the isolates might be from community settings rather than nosocomial settings. Further, there was no apparent difference in prevalence within the sex category. However, in the case of sample type, more than half of the CREc isolates from urine samples harbored *intI1*, which may be a selection bias as majority of the sample were urine. However, studies have always shown that *E. coli* are the most dominant uropathogens^4^ and are more inclined to AMR development.^16^ High empiric use of antibiotics for UTI treatment is the main reason for such inclination of AMR development in uropathogens including *E. coli* in countries like Nepal.^33^

In the antibiotic resistant profile, all the isolates were carbapenem resistant due to selection bias since the study population is CRE isolates. However, the resistant pattern of these CRE isolates towards other classes of antibiotics revealed that integron-positive population showed higher resistance to Penicillins (Amoxicillin, Piperacillin), and Fluoroquinolones (Ciprofloxacin, Levofloxacin, Ofloxacin); followed by Cephalosporins (Cefotaxime, Cefepime, Cefoxitin, Ceftazidime). And, several similar studies done in China reported similar higher antibiotic resistance in integron-positive populations. For instance, in 2022 a study^26^ on clinical *E. coli* isolates revealed a higher resistance to Ampicillin (93%), Ciprofloxacin (80%) and Cotrimoxazole (76%). Likewise, clinical *E. coli* ST131 have been reported to show higher resistance to Cotrimoxazole (95%), Cefotaxime (68%), Ciprofloxacin (68%), Levofloxacin (68%) within integron-positive population^6^. Reportedly, not just *E. coli* but Enterobacteriaceae family as a whole has been reported with higher AMR specifically Ampicillin (100%), Ampicillin-sulbactam (100%), Cefazolin (100%), Levofloxacin (97%) and Ciprofloxacin (96%) within the integron-positive population in China.^25^ Besides China, similar high resistance has been reported with integron-positive population from a hospital of Pakistan^23^ where higher resistance was shown towards Amoxicillin-clavulanic acid (100%), Ceftazidime (94%) and Cefotaxime (94%). Evidently, it is believed that overall antibiotic susceptibility will likely be high in bacteria if they do not have integrons but the frequency of integron negative population of this study is very low, therefore the relation between AMR and integron-population could not be reported with certainty in this study.

In case of Meropenem MIC, the value showed clustering of isolates towards higher MIC and both the MIC_50_ and MIC_90_ had higher value than the breakpoint which questions the current breakpoint efficacy to detect Meropenem resistance. Despite this, the distribution of Meropenem resistance (based on MIC value) within the integron-positive and negative population was statistically significant (p<0.05).

As for carbapenemase production, greater proportion of carbapenemase producers (98.1%) was identified, which is very likely because carbapenemase production is the most common resistance mechanism within CRE strains.^2-3^ And even within the Class 1 integron-positive population, the carbapenemase producing strains were high (n=47/48). Phenotypic testing confirmed all except one isolate to be producing carbapenemases, however, the same isolate was positive for both OXA-23 and OXA-48 and this warrants further genomic investigations to explain why this isolate did not produce carbapenemases in the test result. Further, among the carbapenemase producing isolates (n=51), two isolates were negative for all 6 carbapenemase (NDM-1, OXA-23, OXA-48, VIM, KPC, IMP) encoding genes tested. This genotypic and phenotypic result discrepancy may be due to the fact that the enzyme-producing trait of the CREc isolate is not solely attributed to the six CRE *bla* gene variants (NDM-1, OXA-23, OXA-48, VIM, KPC, or IMP) that were tested in this study. To endorse the carbapenemase producing trait with certainty several genes encoding carbapenemase like CTX-M, SHV, GES, TEM along with their variants which were not explored in this study should be tested using genotypic method. In this study KPC and IMP genes were not detected in any of the isolates, which is in agreement to the findings from a study in India on clinical CREc. ^24^ However, majority of the isolates (79.2%) were positive for NDM-1, followed by OXA-23, OXA-48, and VIM within the integron-positive population. The dominance of NDM-1 is not surprising, given that it is the most common carbapenemase gene that is found in *E. coli* ^24-27^ and; NDM-1 is particularly associated with class 1 integrons gene cassette and IncM plasmids which help them to mobilize.^34^ So the co-occurrence on NDM-1 with *intI1* as well as with other OXA variants is concerning as they can transfer the antibiotic resistant trait by moving between species.^34, 36^ Additionally, the presence of the VIM gene, though only found in three isolates (6.2%) in our study, is interesting as there are only few studies reporting VIM presence in Enterobacteriaceae particularly *Klebsiella pneumoniae, Escherichia coli* and *Enterobacter* spp.^35^ But most importantly, literature suggests that both of these amber class B carbapenemase-encoding genes are associated with integrons and are basically integrated into the gene cassette,^36^ so their 100% prevalence within the integron-positive isolates in this study is relevant. Furthermore, though the OXA-23 and OXA-48 genes were also detected in the integron-positive population but 3 out of 4 integron-negative isolates in this study had OXA variant genes solely. This supports the fact that, these OXA are not commonly integron-associated and are typically encoded in the chromosome,^37^ but their association with plasmids has been reported. Interestingly, both of these Amber class D carbapenemase-encoding *bla* genes (OXA-23 and OXA-48) are typically found in non-Enterobacteriaceae bacteria,^36-37^ and due to their ability to rapidly mutate and expand their spectrum of activity,^36^ their occurrence among *E. coli* is quite noteworthy even in this study. Each of the six CRE gene tested in this study were found discretely (18.8%) as well as in different combination (77.1%) with each other within integron-positive population. The most predominant (13 isolates) combination was NDM-1 and OXA-23, which has already been reported in Nepal but in *Acinetobacter* spp. ^38^ And out of them, 12 isolates were integron positive. So, this combination occurrence in *E. coli* is interesting and might be an indication of probable gene transfer within different species.

Nevertheless, the presence of this variety of carbapenemase combinations within integron-positive bacteria is still concerning, as integrons can provide a platform for AMR gene recombination, creating a novel gene each time and becoming a menace in disease management in Nepal.^11-12^ Clearly, in Nepal, the AMR among the bacteria is the consequence of uncontrolled prescription and over-the-counter use, so unless there is a behavior change in the prescription pattern and controlled use of antibiotic consumption in the Nepalese community, even the newly developed class of antibiotics will eventually develop resistance. Therefore, a proper AMR stewardship program focusing on behavior change along with routine epidemiological surveillance could be pivotal in controlling the AMR problem in Nepal.

To summarize, the class 1 integrons are highly prevalent in multi-drug-resistant *E. coli*, particularly the carbapenemase producing Enterobacterales causing severe infections. And such a high prevalence of carbapenemase producers within class 1 integron-positive *E. coli* possessing resistance to a variety of antibiotic classes is alarming as they can even lead to the creation of pandrug-resistant bacteria. Therefore, the occurrence as well as genomic analysis of class 1 integrons exploring the mechanism should be studied to intercept their evolution and dissemination within the bacterial population.

## Acknowledgements

The authors acknowledge to Mr. Saroj Paudel of Nepalese Farming institute for technical support. We are thankful to all individuals whose samples were used in this study. We also acknowledge Kathmandu Model Hospital for laboratory support.

## Funding

This work was financially supported by NAST research grant (2078/79).

## Transparency declaration

The authors have no conflicts of interest to declare.

## Author Contributions

**ST:** Conceptualization, Funding acquisition, Supervision, Validation, Data analysis, Draft manuscript preparation, Reviewing and editing; **DRJ:** Funding acquisition, Supervision, Validation, Reviewing and editing; **BS, RT, EJ:** Supervision, Reviewing and editing; **MS, YP:** Lab investigation, Reviewing and editing

## Notes

### Competing Interest Statement

The authors have declared no competing interest.

## References

1. World Health Organization. Prioritization of pathogens to guide discovery, research and development of new antibiotics for drug-resistant bacterial infections, including tuberculosis. World Health Organization; 2017.

2. Nordmann P, Poirel L. Epidemiology and diagnostics of carbapenem resistance in gramnegative bacteria. Clinical Infectious Diseases. 2019 Nov 13;69 (Supplement_7):S521–8. 10.1093/cid/ciz824

3. Aurilio C, Sansone P, Barbarisi M et al. Mechanisms of Action of Carbapenem Resistance. Antibiotics. 2022;11(3):1–8.. 10.3390/antibiotics11030421

4. Fluit A, Schmitz FJ. Class 1 integrons, gene cassettes, mobility, and epidemiology. European Journal of Clinical Microbiology and Infectious Diseases. 1999 Dec; 18:761–70.

5. Gillings MR. Integrons: past, present, and future. Microbiology and molecular biology reviews. 2014 Jun;78(2):257–77.10.1128/MMBR.00056-13

6. Huang J, Lan F, Lu Y et al. Characterization of Integrons and Antimicrobial Resistance in Escherichia coli Sequence Type 131 Isolates. Can J Infect Dis Med Microbiol. 2020; Feb 24 2020. 10.1155/2020/3826186

7. Canal N, Meneghetti KL, De Almeida CP et al. Characterization of the variable region in the class 1 integron of antimicrobial-resistant Escherichia coli isolated from surface water. Brazilian J Microbiol. 2016;47(2):337–44. 10.1016/j.bjm.2016.01.015

8. Vásquez-Ponce F, Higuera-Llantén S, Parás-Silva J et al. Genetic characterization of clinically relevant class 1 integrons carried by multidrug resistant bacteria (MDRB) isolated from the gut microbiota of highly antibiotic treated Salmo salar. J Glob Antimicrob Resist. 2022;29:55–62. 10.1016/j.jgar.2022.02.003

9. Firoozeh F, Mahluji Z, Khorshidi A et al. Molecular characterization of class 1, 2 and 3 integrons in clinical multi-drug resistant Klebsiella pneumoniae isolates. Antimicrobial Resistance & Infection Control. 2019 Dec;8:1–7. 10.1186/s13756-019-0509-3

10. Li L, Zhao X. Characterization of the resistance class 1 integrons in Staphylococcus aureus isolates from milk of lactating dairy cattle in Northwestern China. BMC Vet Res. 2018;14(1):1–7. 10.1186/s12917-018-1376-5

11. Pokhrel RH, Thapa B, Kafle R et al. Co-existence of beta-lactamases in clinical isolates of Escherichia coli from Kathmandu, Nepal. BMC Research Notes. 2014 Dec;7:1–5. 10.1186/1756-0500-7-694

12. Tamang MD, Oh JY, Seol SY et al. Emergence of multidrug-resistant Salmonella enterica serovar Typhi associated with a class 1 integron carrying the dfrA7 gene cassette in Nepal. Int J Antimicrob Agents. 2007;30(4):330–5. 10.1016/j.ijantimicag.2007.05.009

13. KC S, Khanal S, Joshi TP et al. Antibiotic resistance determinants among carbapenemase producing bacteria isolated from wastewaters of Kathmandu, Nepal. Environmental Pollution. 2024 Feb 15;343:123155.

14. Thakali O, Malla B, Raya S et al. Prevalence of antibiotic resistance genes in drinking water of the Kathmandu Valley, Nepal. Environmental Challenges. 2022 Apr 1;7:100527.

15. Thakali O, Tandukar S, Brooks JP et al. The occurrence of antibiotic resistance genes in an Urban River in Nepal. Water. 2020 Feb 7;12(2):450.

16. Souque C, Escudero JA, Maclean RC. Integron activity accelerates the evolution of antibiotic resistance. Elife. 2021;10:1–47. 10.7554/eLife.62474

17. Clinical and Laboratory Standards Institute. Performance Standards for Antimicrobial Susceptibility Testing: Thirty-First Edition M100-S31. CLSI, Wayne, PA, USA, 2022

18. Cheesbrough M. District laboratory practice in tropical countries. Cambridge University Press; 2010.

19. Hartas J, Hibble M, Sriprakash KS. Simplification of a locus-specific DNA typing method (Vir typing) for Streptococcus pyogenes. Journal of clinical microbiology. 1998 May 1;36(5):1428–9. 10.1128/jcm.36.5.1428-1429.1998

20. Colello R, Krüger A, Di Conza J et al. Antimicrobial resistance in class 1 integron-positive shiga toxin-producing Escherichia coli isolated from cattle, pigs, food and farm environment. Microorganisms. 2018;6(4):2–9. 10.3390/microorganisms6040099

21. Mahmoud NE, Altayb HN, Gurashi RM. Detection of carbapenem-resistant genes in Escherichia coli isolated from drinking water in Khartoum, Sudan. J Environ Public Health. 2020;2020.

22. Tawfick MM, Alshareef WA, Bendary HA et al. The emergence of carbapenemase bla NDM genotype among carbapenem-resistant Enterobacteriaceae isolates from Egyptian cancer patients. Eur J Clin Microbiol Infect Dis. 2020;39(7):1251–9. 10.1007/s10096-020-03839-2

23. Al-Hammadi MA, Al-Shamahy HA, Ali AQ et al. Class 1 integrons in clinical multi drug resistance E. coli, sana’a hospitals, Yemen. Pakistan J Biol Sci. 2020;23(3):231–9. 10.3923/pjbs.2020.231.239

24. Joshi DN, Shenoy B, Mv B et al. Prevalence of Carbapenem-Resistant Enterobacteriaceae and the Genes Responsible for Carbapenemase Production in a Tertiary Care Hospital in South India. Eur Med J. 2023;66(May 2023). 10.33590/emj/10300425

25. Wang T, Zhu Y, Zhu W et al. Molecular characterization of class 1 integrons in carbapenem-resistant Enterobacterales isolates. Microb Pathog. 2023;177(February):106051. Available from: 10.1016/j.micpath.2023.106051

26. Li W, Ma J, Sun X et al. Antimicrobial Resistance and Molecular Characterization of Gene Cassettes from Class 1 Integrons in Escherichia coli Strains. Microbial Drug Resistance. 2022 Apr 1;28(4):413–8. 10.1089/mdr.2021.0172

27. Leverstein-Van Hall MA, Blok HEM et al. Multidrug resistance among Enterobacteriaceae is strongly associated with the presence of integrons and is independent of species or isolate origin. J Infect Dis. 2003;187(2):251–9. 10.1086/345880

28. Barzegar S, Arzanlou M, Teimourpour A et al. Prevalence of the Integrons and ESBL Genes in Multidrug-Resistant Strains of Escherichia coli Isolated from Urinary Tract Infections, Ardabil, Iran. Iran J Med Microbiol. 2022;16(1):56–65.

29. Kang HY, Jeong YS, Oh JY et al. Characterization of antimicrobial resistance and class 1 integrons found in Escherichia coli isolates from humans and animals in Korea. J Antimicrob Chemother. 2005;55(5):639–44. 10.1093/jac/dki076

30. Seo KW, Lee YJ. Prevalence and Characterization of Plasmid Mediated Quinolone Resistance Genes and Class 1 Integrons among Multidrug-Resistant Escherichia coli Isolates from Chicken Meat. J Appl Poult Res. 2019;28(3):761–70. Available from: 10.3382/japr/pfz016

31. Sütterlin S, Bray JE, Maiden MCJ et al. Distribution of class 1 integrons in historic and contemporary collections of human pathogenic Escherichia coli. PLoS One. 2020;15(6):1–13. 10.1371/journal.pone.0233315

32. Huang L, Huang C, Yan Y et al. Urinary Tract Infection Etiological Profiles and Antibiotic Resistance Patterns Varied Among Different Age Categories: A Retrospective Study From a Tertiary General Hospital During a 12-Year Period. Front Microbiol. 2022;12(January):1–10. 10.3389/fmicb.2021.813145

33. Parajuli NP, Maharjan P, Parajuli H et al. High rates of multidrug resistance among uropathogenic Escherichia coli in children and analyses of ESBL producers from Nepal. Antimicrob Resist Infect Control. 2017;6(1):1–7. Available from: 10.1186/s13756-016-0168-6.

34. Lopez-Diaz M, Ellaby N, Turton J et al. NDM-1 carbapenemase resistance gene vehicles emergent on distinct plasmid backbones from the IncL/M family. Journal of Antimicrobial Chemotherapy. 2022 Mar 2;77(3):620–4. 10.1093/jac/dkab466

35. Matsumura Y, Peirano G, Devinney R et al. Genomic epidemiology of global VIM-producing Enterobacteriaceae. J Antimicrob Chemother. 2017;72(8):2249–58. 10.1093/jac/dkx148

36. Diene SM, Rolain JM. Carbapenemase genes and genetic platforms in Gram-negative bacilli: Enterobacteriaceae, Pseudomonas and Acinetobacter species. Clinical Microbiology and Infection. 2014 Sep 1;20(9):831–8.

37. Walther-Rasmussen J, Høiby N. OXA-type carbapenemases. J Antimicrob Chemother. 2006;57(3):373–83. 10.1093/jac/dki482

38. Joshi PR, Acharya M, Kakshapati T et al. Co-existence of bla OXA-23 and bla NDM-1 genes of Acinetobacter baumannii isolated from Nepal: antimicrobial resistance and clinical significance. Antimicrobial Resistance & Infection Control. 2017 Dec;6(1):1–7.

