## Supplementary Figures for "Prevalence of Class 1 integron in disease-causing carbapenem-resistant *Escherichia coli* in a tertiary hospital in Kathmandu, Nepal"

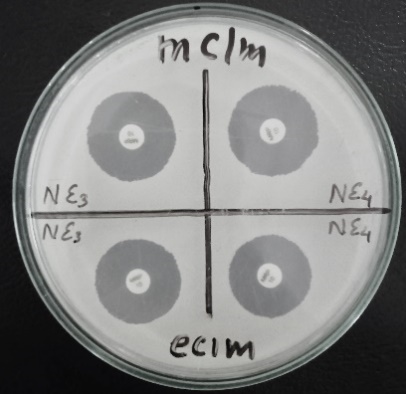

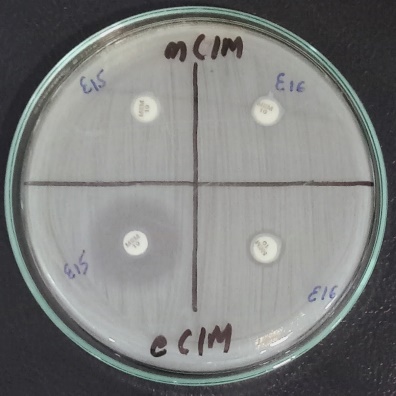

(A) (B)

***Supplementary Figure F1*:** *Zone of inhibition shown during eCIM and mCIM test: carbapenemase negative (A), MBL producer (B, left side) and SBL producer (B, right side)*

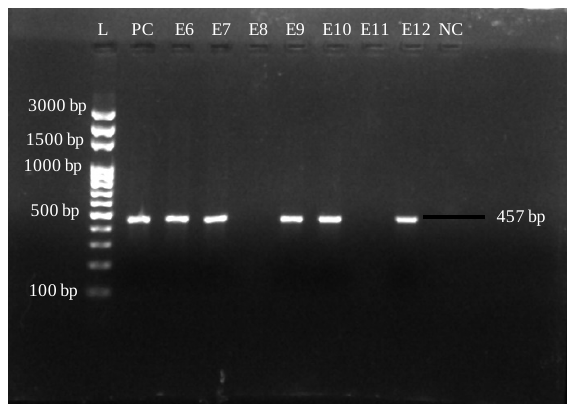

***Supplementary Figure F2****: Gel visuals showing the presence of class 1 Integron gene in E6, E7, E9, E10 and E12 sample of band length 457bp while no band in E8 and E11 sample indicating absence of gene. (L is ladder, NC is negative control, PC is positive control and E is sample).*

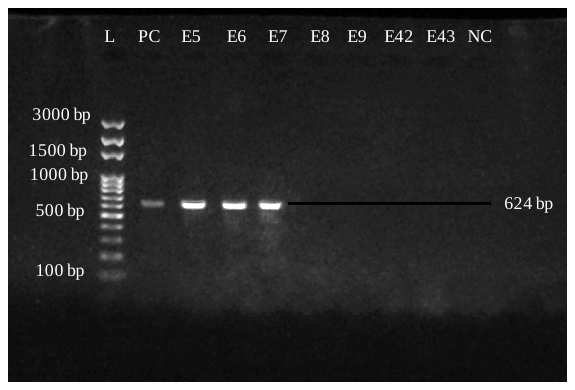

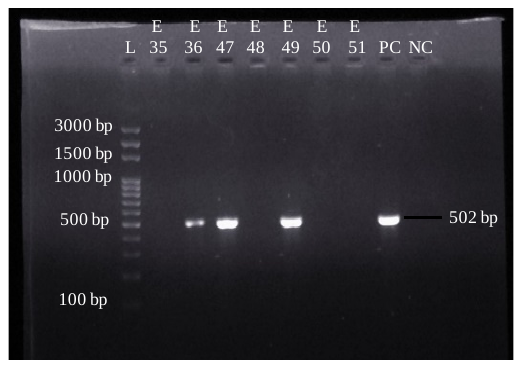

| (A)  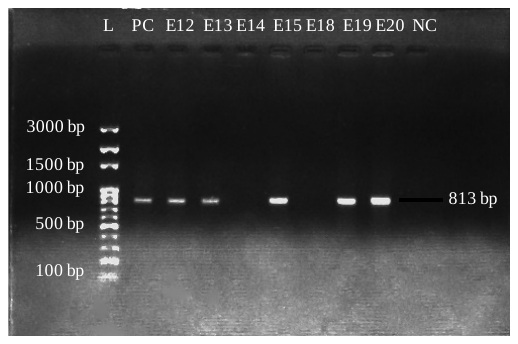 | (B)  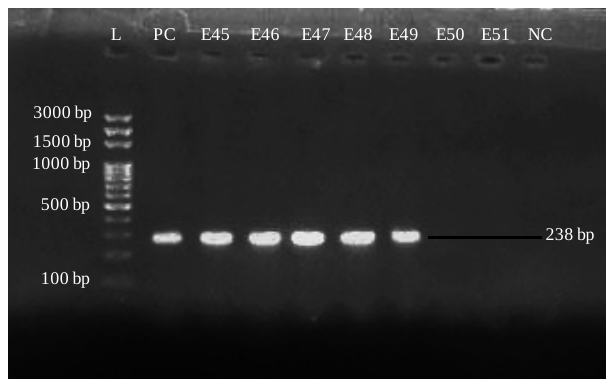  1500 bp  238 bp |
| --- | --- |
| (C) | (D) |

***Supplementary Figure F3***: *Gel bands showing the presence of NDM-1 (A), VIM (B), OXA-23 (C) and OXA-48 (D).*
